# Dancing bees evaluate agricultural forage resources as inferior to central urban land

**DOI:** 10.1101/2019.12.19.882076

**Authors:** Ash E. Samuelson, Roger Schürch, Ellouise Leadbeater

**Affiliations:** School of Biological Sciences, Royal Holloway University of London, Egham, United Kingdom; Department of Entomology, Virginia Tech, Blacksburg, VA, USA

## Abstract

Recent evidence suggests that flower-rich areas within cities could play an important role in pollinator conservation, but direct comparison of agricultural and urban areas has proved challenging to perform over large scales. Here we use the waggle dances of honeybees (*Apis mellifera* L.) to evaluate floral resource availability over the entire season at deeply urban or agricultural sites. Through analysis of 3378 dances that were performed over two years by 20 colonies in SE England, we show that foraging trip distance is consistently lower at urban sites across the entire season, implying a higher availability of forage in heavily urbanized areas. Urban bees also collected nectar with a higher mean sugar content. From the self-reported perspective of a generalist pollinator, the modern agricultural landscapes that we studied provided insufficient and transient resources relative to heavily urbanised areas, which may represent important refuges within an impoverished landscape.

## Introduction

The most pressing threat facing bee populations worldwide is habitat loss and fragmentation, mediated by agricultural intensification over the last century ^1,2^. In combination with challenges posed by widespread pesticide use^3^ and emerging parasites and disease^4^, this extensive conversion of flower-rich habitat to land that is often nutritionally barren from the bee’s perspective has been strongly implicated as a driver of wild bee declines^2,5,6^. Within this context, growing evidence to suggest that flora-rich patches within cities and towns may support diverse bee populations^7,8^ and that social bee colonies in urban areas outperform their agricultural counterparts^9,10^ has led to suggestions that cities might offer important refuges in an impoverished agricultural landscape. Although surrounded by large expanses of impermeable man-made surface, urban parks, gardens and allotments can offer high floral abundance and diversity across the season^8,11^. However, directly comparing forage resources for bees in urban and agricultural environments through traditional surveying presents a major challenge of both access and scale that has thus far proved hard to overcome.

Here, we capitalize upon the unique communication behaviour of a generalist pollinator to compare the floral resources available to bees in urban and agricultural environments at the landscape scale. Honeybees collect food from a broad range of floral resources across a vast foraging range (up to a 10km radius^12^), visiting many species also visited by wild bee communities^13^. Successful foragers communicate locations of profitable resources to their nestmates, by performing a figure-of-eight “waggle dance” on the comb that encodes the distance to the resource (in the duration of the “waggle” run) and the angle from the sun’s azimuth (in the angle of the dance relative to gravity^14^). By decoding these dances, it is possible to obtain filtered real-time information about the forage sites that have been found by the hive’s workforce^15^ that is relatively less affected by proximity to local hotspots than traditional surveying, with no access limitations (a key hurdle in surveying urban areas). Because honeybee colonies are economical foraging entities that are unlikely to focus upon distant resources when near ones of similar quality are available, the distance of these sites from the hive acts as a proxy for forage availability^16–18^, while the quality of forage can be independently verified by non-destructive assay of the sucrose content of forager-collected nectar^16^. Thus, honeybee colonies can be used to survey landscapes comprehensively, giving a real-time picture of current forage availability and quality within their foraging range.

In the most geographically extensive waggle dance study to date, we decoded 2827 dances from twenty observation hives placed at either the urban or the agricultural extremes of an urbanization gradient in SE England (Fig. 1), recorded fortnightly over 24 weeks from April-September 2017. We also analysed 551 dances from a subset of these hives in 2016, to investigate consistency across years. We compared foraging distance, as a proxy for forage availability, between the two landscape types, alongside analysis of nectar quality (sugar concentration) to investigate whether differences in foraging distance might be energetically compensated for by differences in quality. By mapping dance distributions onto land-use maps (Figs. 2 & S2), we also investigated the importance of specific land-use types within the urban and agricultural sites across the season.

**Figure 1.**
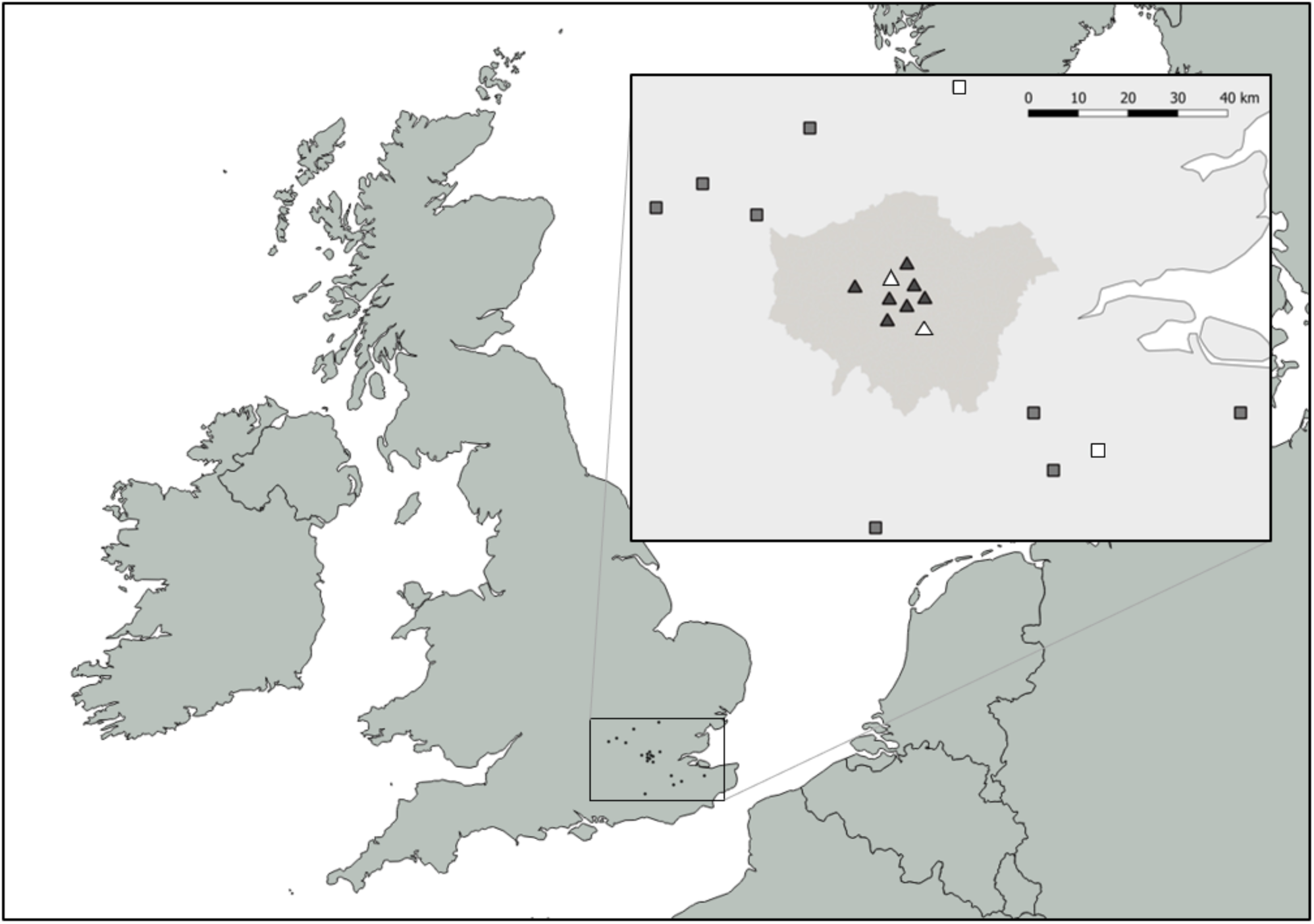
Location of ten urban (triangles) and ten agricultural (squares) observation hives in SE England. In the inset map the Greater London area is indicated by dark grey shading. Hives were located in the highly urbanised centre of London and the agricultural areas around London to represent extremes of an urbanisation gradient and were a minimum of 5000m apart to minimise overlap of foraging ranges. In 2017, dances were recorded from all hives; the 2016 subset is indicated by unfilled symbols.

**Figure 2.**
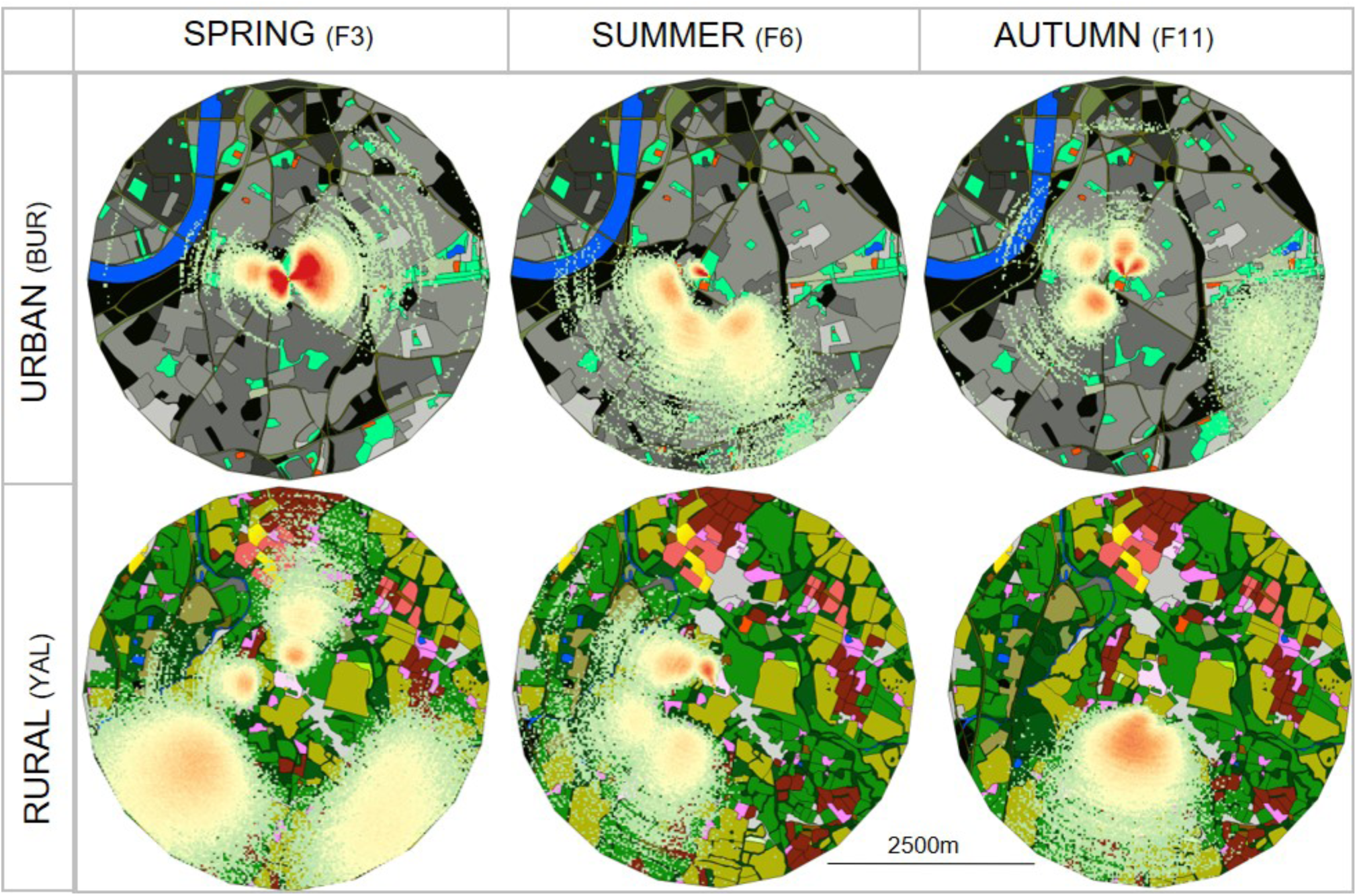
Example waggle dance plots from one urban site (BUR) and one agricultural site (YAL). Each circle shows the dances recorded on a single filming period (up to 3 hours) during spring (fortnight 3), summer (fortnight 6) and autumn (fortnight 11). Waggle dances are displayed as probability heatmaps generated from 1000 simulations of each dance allowing incorporation of variability in distance and angle communication ^29^. Dance plots are overlaid on GIS land-use maps (radius 2500m) produced for land-use preference analysis. For waggle dance plots for all 183 site-fortnight combinations see Fig S1.

## Results

### Distance to forage and nectar quality

Across the whole season, we found a strong overall effect of land-use on median hive waggle run duration, implying that bees flew further to find forage in agricultural sites (median translated foraging distance: 1108m; maximum: 8599m) than urban sites (median 708m; maximum: 9523m; Fig. 3a; ΔAICc to null model: 3.06, Table S1a; land-use parameter estimate [95% CIs]: 1.180 [0.168 to 2.192], Table S2a). This difference was greatest in the spring, when urban bees travelled shorter distances to find food than they did in the summer months (Fig 3a; significance of separate urban and agricultural smooth terms in GAMM: *p* <0.001; Table S2a). In agricultural sites duration did not follow a strong seasonal pattern, most likely because the timings of peaks in availability (consistent with mass-flowering crops) varied between individual agricultural sites (Fig. 4; significantly higher standard deviation in median log-transformed durations across sites in agricultural hives than urban; ΔAICc to null model: 2.63; land-use parameter estimate: 0.122 [0.019 to 0.224]). However, even in the late season when urban bees flew further to find food than in the spring, agricultural bees still had to travel even longer distances (Fig 3a). There was no effect of temperature, (parameter estimate: 0.011 [-0.019 to 0.042]), filming period (AM or PM; parameter estimate: −0.033 [−0.138 to 0.072]; Table S2a), or dance decoder identity (not included in final 95% confidence model set; ΔAICc from best model to model containing decoder variable: 2.31; Table S1a) on waggle run duration.

**Figure 3.**
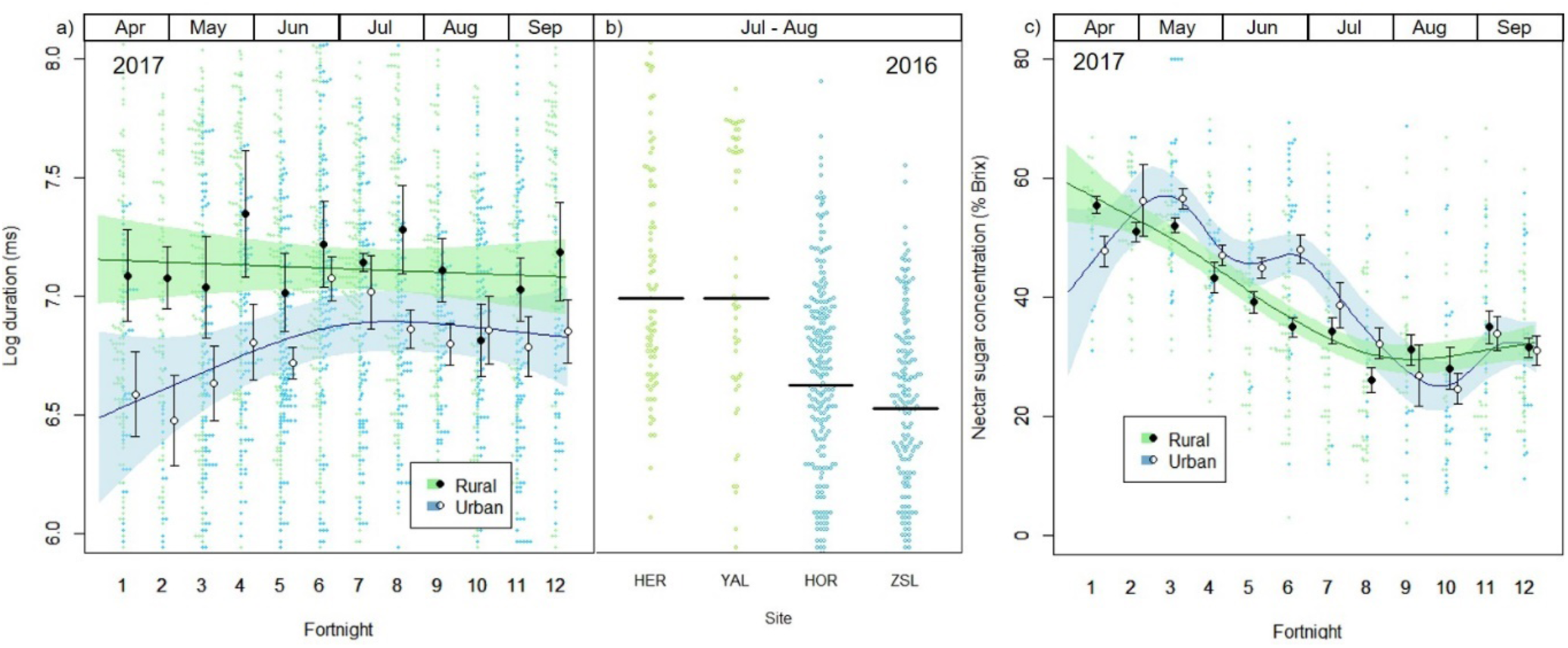
a) Median log-transformed waggle run duration ± SE for urban (open circles) and agricultural (filled circles) colonies over twelve fortnightly timepoints between April and September 2017. Waggle run duration across all sites and fortnights ranged from 0.12 to 12.44 sec, which translates to a minimum distance of 56m and a maximum distance of 9523m, with the median distance 897m. Lines are fitted from GAMs allowing a non-linear relationship between fortnight and duration, with shaded areas indicating 95% CIs. Raw data are shown in the background beeswarm plot (blue: urban, green: agricultural). b) Beeswarm plot of log-transformed waggle run durations for dances recorded over eight weeks in July and August 2016 at a subset of two urban (blue) and two agricultural (green) sites. Black lines indicate median values. c) Median nectar sugar concentration (% Brix) ± SE from nectar collected from returning foragers immediately after waggle dance data collection in 2017 (see (a) for information on plot features).

**Figure 4.**
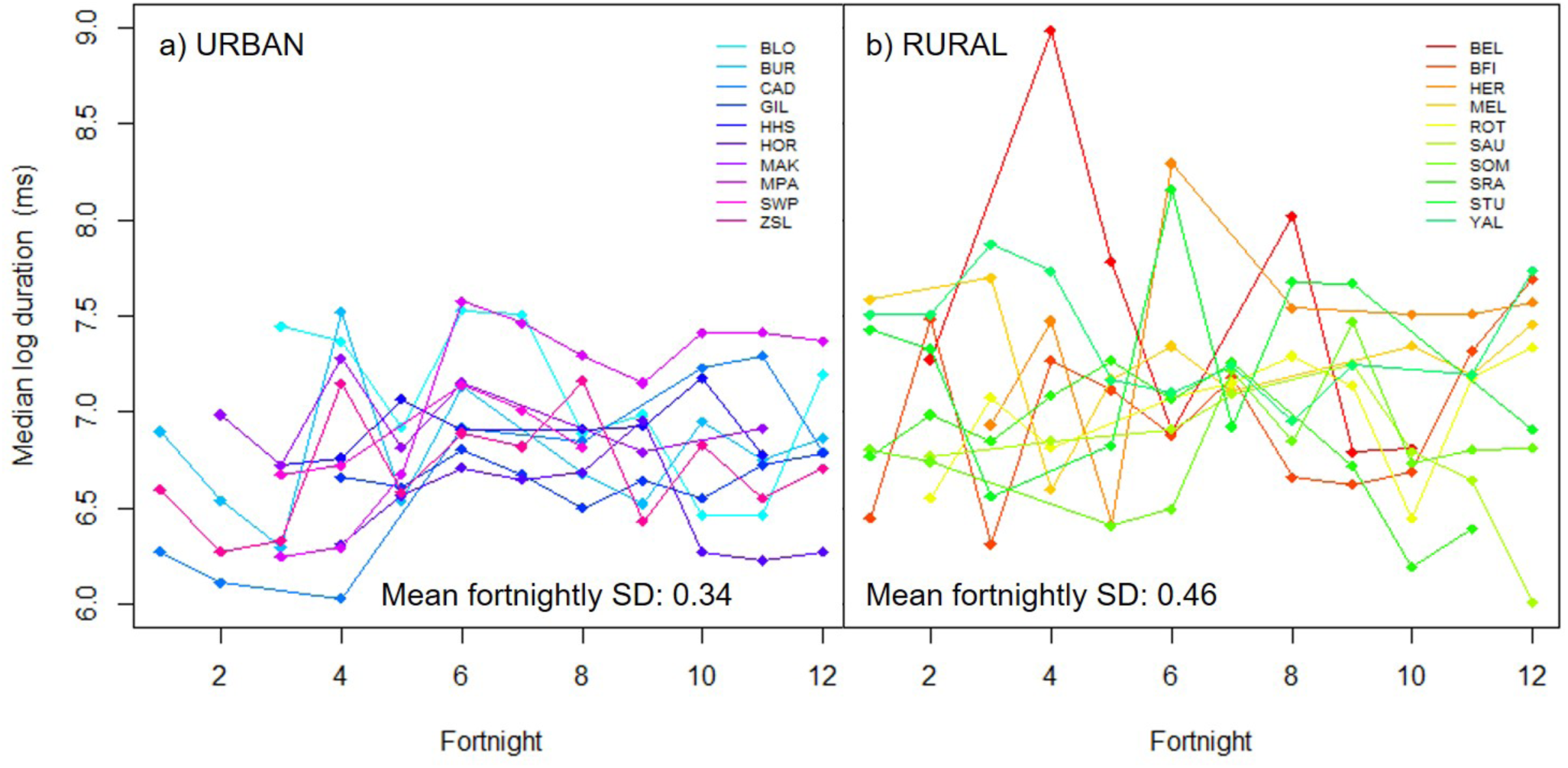
Median log-transformed waggle run durations for individual a) urban and b) agricultural sites over twelve fortnightly timepoints, showing greater variation in foraging distance across agricultural sites than urban sites (mean standard deviation is inset). Variation in peaks of forage availability in agricultural sites suggest reliance on a few ephemeral crop resources, the identity of which differ between sites. In contrast urban sites follow a consistent seasonal pattern suggesting exploitation of a larger number of resources.

Data from 551 dances decoded from a subset of hives (two urban and two agricultural) collected in 2016 produced similar results (Fig. 3b and S2), with a strong effect of land-use on duration (ΔAICc to null model: 5.94, Table S1c; land-use parameter estimate: −0.836 [−1.664 to −0.007], Table S2c). Translated to foraging distance, the urban median was 475m (mean: 540m; maximum: 2517m) and the agricultural median was 927m (mean: 1288m; maximum: 3979m), indicating consistency across years.

Analysis of the nectar collected by returning foragers showed that longer flights in agricultural landscapes were not compensated for by collection of high-quality resources (Fig. 3c). Land-use and its interaction with season affected nectar quality (sugar concentration), with overall higher nectar quality in urban land (urban mean: 41.38(±0.99) °Brix) than agricultural (agricultural mean: 38.02(±0.73) °Brix; ΔAICc to null model: 279.91, Table S1b; land-use parameter estimate: 22.020 [7.547 to 36.493], Table S2b). In both land-use types, nectar quality declined over the season, although this decline was less smooth in urban land (Fig. 3c).

### Land-use preference

Analysis of preference for small-scale land-use types within the wider urban and agricultural landscapes highlighted reliance on residential gardens in urban areas and mass-flowering crops in agricultural areas (Fig. 5). Specifically, bees in urban areas showed a clear preference for sparse residential land (discontinuous development typified by large total garden area) across the whole season (spring odds ratio (OR) [95% CIs]: 5.4 [3.0 to 10.0]; summer: 4.6 [2.9 to 7.2]; autumn: 4.2 [2.6 to 6.8]) and a relatively smaller yet still significant preference for dense residential land (discontinuous development with a high ratio of impervious surface to gardens; spring OR: 2.4 [1.3 to 4.3]; summer: 3.4 [2.1 to 5.8]; autumn: 3.3 [1.9 to 5.5]). In agricultural areas in spring, bees showed a strong preference for oilseed rape fields (OR: 24.8 [13.0 to 46.0]).

**Figure 5.**
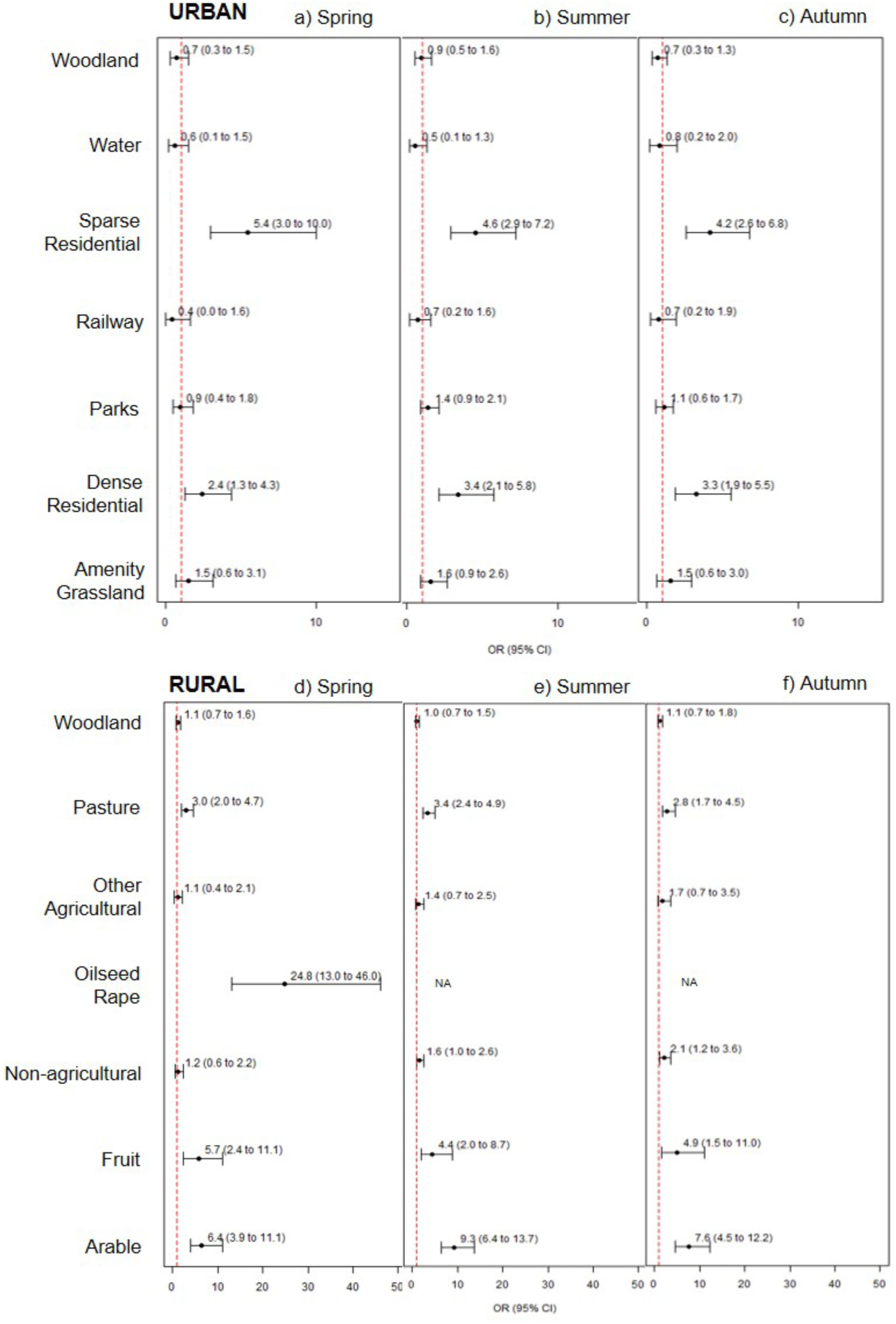
Adjusted odds ratios (± 95% CIs) for visitation to land-use types within urban (a-c) and agricultural (d-f) landscapes derived from simulated foraging visits using 2017 waggle dance data. An odds ratio of 1 is indicated by the red dashed line. Error bars that do not cross 1 indicate significant differences from the baseline land-use type (“Continuous NUrban” in Urban and “Built-Up” in Agricultural). Data from fortnights 1-4 (spring), 5-8 (summer) and 9-12 (autumn) are pooled.

## Discussion

Despite being surrounded by land dominated by man-made surfaces, the urban honeybees in our study had to travel significantly less far to find food throughout the season than those in agricultural areas, suggesting that urban areas provide consistently more available forage. In large portions of Western Europe and North America, agricultural intensification has led to the removal of hedgerows and wildflowers^2^, leaving ephemeral mass-flowering crops as the major pollen and nectar resource for those pollinators that can exploit them^19^. In our study, the peaks in agricultural forage availability indeed reflected visits to these crops, but the troughs that preceded or followed them led to overall increased foraging distances that were not recompensed by increased nectar sugar content; indeed, nectar sugar concentration was lower overall in agricultural areas. Our findings demonstrate the value of urban forage resources within these intensified agricultural landscapes for *Apis mellifera*-a species that is both an important pollinator and a key model for understanding the foraging dynamics of social bees.

*A. mellifera* is the only *Apis* species that is present in the western European environment that we studied, but it is likely that aspects of these findings are generalizable to other generalist social bees^15^, including several *Bombus* spp. Both managed *Apis* and wild *Bombus* are critically important ecosystem service providers within these landscapes^20,21^. The picture is less clear for more specialist and/or solitary bee species, for which the presence of particular flower species as larval food sources may well be key^5^. However, recent work^8,22^ has identified residential areas within cities as “hotspots” for a wide range of pollinators, including solitary bees. While food availability is unlikely to be the only determinant of bee species richness and abundance within cities, because nesting site availability will also be key, it seems clear that these residential areas offer a precious foraging resource for a broad pollinator community within urban landscapes. Although generalization to the species level should be undertaken with caution, our direct comparison of urban and agricultural land shows that such areas may represent important refuges for bees within an inhospitable agricultural landscape.

The availability of urban green space varies both between and within cities^23^, and our results from urban sites that were specifically chosen to represent the extreme, central end of an urbanization spectrum (Fig. 1) most likely under-estimate the floral resources available in more suburban areas. On a wider scale, given that northern European cities such as London typically contain relatively high green-space availability^23^, our findings support that urban planning that prioritizes residential green space could increase the value of urban areas for bees in cites that are less garden-rich. Nonetheless, urban land remains a small percentage of total land cover and these islands of abundant forage may be insufficient to support bee populations across a landscape dominated by intensive agriculture. As such, in the long term, conservation efforts should be primarily directed towards increasing non-crop floral provision in agricultural areas, such as wildflower strips^24^, to increase consistency of forage availability across the season and to minimise reliance on small numbers of ephemeral flowering crops. Our study harnesses the unique habitat surveying capability of the honeybee waggle dance to demonstrate that cities can provide islands of forage within relatively barren agricultural land, but since crop plants are located outside of cities, redressing this balance through improved floral provision on agricultural land will be critical to healthy ecosystem service provision.

## Methods

### Experimental Design

We selected ten highly urban (central London) and ten agricultural sites in SE England that either had existing observation hives (n=10) or an existing apiary where it was possible to situate an observation hive for the duration of the experiment (n=10; Fig. 1; see “Land use preference analysis” for site classification methods). Sites were located at least 5000m apart to minimise overlap of foraging ranges and were selected to be representative of the extremes of the urbanisation spectrum in the region. In 2017, dances from each site were videoed once every two weeks for 24 weeks between April and September (a total of 12 recordings from each site; two sites visited each day between 8:00-12:00 (“AM”) and 12:00-17:00 (“PM”) respectively). Duration of the central waggle run of a dance correlates linearly with the distance to the foraging site that a bee has visited^14,25^, so we compared the median waggle dance duration (time to perform the central waggle run) in each video (183 site-fortnight combinations) in order to assess forage availability in urban and rural sites across the season (see “Data Analysis”). We also compared the concentration of nectar in the crop of ten returning foragers captured from each hive after each video recording session, in order to evaluate differences in forage quality. The same procedure was followed for two months (July-August) in 2016 at four of the sites (two urban and two agricultural), to investigate whether results were consistent across years.

### Honeybee colonies

An observation hive containing a honeybee colony with 3 to 8 occupied frames of workers, brood and a queen was located at each site. At sites that did not have existing observation hives we used standard three frame hives (two shallow and one deep; Thorne, Windsor, UK) situated in plastic storage sheds (130 × 74 × 110 cm; Keter, Birmingham, UK) with access to the outside through a clear PVC tube (25mm diameter; RS Components, Corby, UK). Existing hives (n=10/20; 6 urban and 4 agricultural) were left *in situ*. Colonies were not supplied with extra food unless they were temporarily at risk of starvation (no nectar in storage cells or poor weather), when they received supplementary sugar syrup (50° Brix) in a gravity feeder at the top of the hive. If colonies failed they were replaced immediately with a new nucleus (n=5 early-season failures). Colonies were checked every few days by apiary managers to ensure adequate stores were available and inspected thoroughly every two weeks. Swarm control (removal of a brood frame and/or queen cells) was carried out between April and July if swarm risk was detected; no colony swarmed during the experiment. At the end of the experiment colonies were merged with others at the same apiary or transferred to nucleus hives at the university.

### Data collection

Visit days for video recording of hives alternated between urban and agricultural pairs of sites. The order of visits was kept approximately consistent throughout the experiment (weather permitting) and the period (AM or PM) of the visit to each hive alternated each fortnight. Sites were visited only on sunny, warm (>12°C) and calm (wind speed <15km/h) days to ensure bees were foraging. Site data recorded included weather and colony strength (number of frames covered with bees to nearest 0.5). Temperature data (daily mean) was taken from the London Heathrow weather station (wunderground.com). In total, we recorded 2827 waggle dances (1428 urban and 1399 agricultural) in 183 site-fortnight combinations in 2017, and 32 site-week combinations in 2016.

On each visit, two hours of waggle dance data were recorded by training a camcorder (Canon Legria HF R606, Amstelveen, NL) onto the dancefloor (the location where most dance activity is seen; *30*). Plumb lines to provide a reference for gravity and a radio-controlled clock were attached to the glass in the field of view. At the end of filming we collected nectar quality data by blocking the entrance to the hive and collecting ten returning foragers not carrying pollen^16^. Following anaesthesia in a cool bag containing ice blocks, we stimulated regurgitation by massaging the bees’ abdomens with forceps. Using a microcapillary tube, crop contents were transferred to a 0-80° Brix refractometer (Kern, Balingen, Germany) to measure sugar concentration.

### Waggle dance decoding

We decoded up to 40 (mean = 15.5) dances per video-recorded session (QuickTime 7.1; frame-by-frame playback at 25fps) following methods outlined by Couvillon *et al*.^26^. Briefly, we decoded four waggle runs for each dance, excluding the first and last runs which typically exhibit more variation than middle runs. For each run we recorded the angle from vertical, and calculated the angular displacement from North of the advertised resource by adding the angle to the sun’s azimuth at the time of the dance (http://www.esrl.noaa.gov/gmd/grad/solcalc/). We measured run duration by recording the first frame in which the bee started vibrating its body and the first frame after the vibration had finished. When a run was interrupted (e.g. by colliding with another bee), that run and the next were skipped. When high dance activity resulted in difficulty keeping track of individual dancers over time, each dance at a single timepoint was decoded and the video skipped forward six minutes to avoid recording the same dance twice. To convert waggle run durations to foraging trip distances we used a universal calibration derived from waggle dance data from multiple honeybee populations worldwide (doi:10.7294/tnm4-9123) which relates waggle run duration to distance using a linear equation and incorporates variation in distance communication^25^.

### Statistical analysis

To analyse waggle run duration (as a proxy for foraging distance) and nectar quality we built GAMMs allowing for a non-linear effect of our time variable, fortnight, on the response, and including site as a random effect. To incorporate an interaction between land-use and fortnight we allowed separate smoothers. For waggle run duration, the response variable was log-transformed median run duration per video. We combined data within videos (representing a single sample session for each hive) because although data collection was designed to minimise the possibility of recording the same bee twice (see *Waggle dance decoding)* it is not possible to identify individual foragers and so we cannot rule out the possibility of a small proportion of dances being carried out by previously recorded individuals, which would cause an analysis of the raw data to violate the assumption of independence. A sensitivity analysis of the raw data resulted in the same results as our main analysis that used video medians. Due to concurvity (non-linear collinearity) between fortnight and temperature, we used sequential regression^27^ to generate the variable residual temperature by regressing temperature against fortnight using a GAM and extracting the residuals to produce a variable containing the variation in temperature not explained by fortnight. “Decoder” was included to test for an effect of which researcher decoded the dances, and was split into the lead researcher (69% of dances) and trained research assistants (n=9, 31% of dances). The full model for waggle run duration (2017 data) contained the covariates land-use, fortnight, residual temperature, filming period (AM or PM) and decoder. Because the 2016 dataset was smaller (n=551 dances) we pooled dances across the study period and instead of using median durations we accounted for non-independence of dances by including recording session as a random effect. Nectar quality (sugar content, °Brix) was analysed with the covariates land-use, fortnight, residual temperature, period (AM or PM) and colony strength. We excluded zero values (2.6% of samples) as these indicate bees that were collecting water^16^. Colony strength (bee-covered surface) was calculated by multiplying the number of frames covered with bees by the surface area of the relevant frames, depending on hive type (National or Commercial) and frame size (Deep or Shallow).

Model selection was carried out using a “full subset” approach with a set of models containing all combinations of covariates plus a null model containing the intercept and random effect. We selected the model with the lowest AICc (Akaike’s Information Criterion corrected for small sample sizes) as the best fitting model(s). Where one or more models were within 2 AIC units of the best model, we performed model averaging on the best model set. Final models were validated graphically to assess fit and check that assumptions had been met, and examined for spatial autocorrelation by using a Moran’s I test on the residuals and graphically assessing the spatial pattern of residuals.

### Land-use preference analysis

We classified the land at a 2500m radius (incorporating the 95^th^ percentile of recorded dances) around each site using QGIS v3.0.2 following methods outlined in ^28^. Briefly, we generated land-use maps (Figure S1) by drawing polygons around habitat patches on a satellite imagery (Bing Maps) base layer and classifying these patches into 33 land-use categories. To ensure that oilseed rape fields (OSR), which may not be detected by satellite imagery, were included in our mapping we additionally performed aerial drone surveys (DJI Phantom 4; DJI, Shenzhen, China; 360° recording from 120m above the hive) at each agricultural site during May (the OSR bloom period).

The strong difference found in foraging distance between urban and agricultural hives led us to investigate which of the land-use types within these differing landscapes received most attention by foraging bees during different seasons (spring: April-May, summer: June-July, autumn: August-September). For each site, we produced a land-use raster of radius 2500m (incorporating the 95^th^ percentile of recorded dances) and resolution 25m. The raster separated land-use patches into broad categories selected for ecological relevance to pollinator use of the landscape ^28^, combined from land-use classes in our initial GIS classification. In agricultural landscapes the categories were: built-up, non-agricultural, woodland, arable, pasture, fruit, oilseed rape and other agricultural; in urban landscapes the categories were: continuous urban, dense residential, sparse residential, parks, amenity grassland, railway, woodland and water (including riverbanks).

For a single site-season combination, we simulated a single foraging location for each recorded dance (mean n=47.86 ±SE 3.22), following methods outlined in^29^ to incorporate variability in angle and distance communication. Each land-use patch was recorded as visited or not visited by one or more of the simulated foraging visits, along with its land-use type, area and distance of nearest edge to the hive. This was repeated for each site in a landscape-season combination, e.g. spring data for all sites in urban landscapes. This generated a data frame containing the variables land-use patch ID, land-use type, visited (0/1), distance from hive, site ID. A binomial GLMM (logit link function) was constructed with visited as the response and fixed effects of land-use and distance (allowing for an inverse relationship between distance and visit probability; *37*) and a random effect of site ID. The adjusted odds ratios (AORs) for visitation of each land-use type relative to the baseline type (selected as the most urban land-use type: “continuous urban” in urban landscapes, “built-up” in agricultural landscape), corrected for distance to the hive, were extracted from the model and stored. This procedure was simulated 1000 times, so that foraging locations reflected the distribution of probabilities defined by the calibration described above. Each iteration generated AORs from the model; the median AOR and 95% confidence intervals (CIs) were extracted. The entire procedure was repeated for each of the six landscape-season combinations. All analyses were conducted in R version 3.2.1 using packages *MuMIn* ^30^, *circular* ^31^, *ascii* ^32^, *raster* ^33^, *sp* ^34^, *rgdal* ^35^, *rgeos* ^36^, *FSA* ^37^, *lme4* ^38^, *beeswarm* ^39^ and *Hmisc* ^40^. Raw data are archived in Dryad (entry doi: xx.xxxx/dryad.xxxx).

## Acknowledgements

This research was supported by the ICL-RHUL BBSRC DTP BB/M011178/1, European Research Council Starting Grant BeeDanceGap 638873, and donations from High Wycombe Beekeepers’ Association and Essex Beekeepers’ Association. We would like to thank Glenn Ahearn, Mehmet Akiner, Sharon Bassey, Peter Buckoke, John Chapple, Terry Clare, Luke Dixon, Melvyn Essen, Bill Fisher, Clive Hill, the Hive Honey Shop, the Horniman Museum & Gardens, James Makinson, Mark Patterson, Sarah & Vincent Rapley, Simon Rice, Louisa Roscoe, Barnaby Shaw, Sarah Turner, Yalding Beekeepers’ Association and ZSL for providing access to and managing observation hives. We are grateful to Huw Fox, Harriet Hall, Jagpreet Hayre, Will Howes, Liberty John, Hana Montague, Michael Sealy, Lucy Tilly-May, Vicky Tubman and Vivitsha Zala for decoding waggle dances, Mark Brown and Rich Gill for advice, Graham Stone for comments on a draft of the manuscript and Lawrence Watson for providing the drone.

## Author Contributions

A.E.S. and E.L. conceived the initial idea and designed the experiment; A.E.S. performed the experiment; A.E.S. & R.S. performed the statistical analyses; A.E.S. wrote the initial manuscript draft and E.L., R.S., and A.E.S. provided the final edit.

## References

1. Potts, S. G. et al. Global pollinator declines: trends, impacts and drivers. Trends Ecol. Evol. 25, 345–353 (2010).

2. Baude, M. et al. Historical nectar assessment reveals the fall and rise of floral resources in Britain. Nature 530, 85–88 (2016).

3. Wood, T. J. & Goulson, D. The environmental risks of neonicotinoid pesticides: a review of the evidence post 2013. Environ. Sci. Pollut. Res. 24, 17285–17325 (2017).

4. Fürst, M. A., McMahon, D. P., Osborne, J. L., Paxton, R. J. & Brown, M. J. F. Disease associations between honeybees and bumblebees as a threat to wild pollinators. Nature 506, 364–366 (2014).

5. Biesmeijer, J. C. et al. Parallel Declines in Pollinators and Insect-Pollinated Plants in Britain and the Netherlands. Science (80-.). 313, 351 LP–354 (2006).

6. Carvell, C. et al. Declines in forage availability for bumblebees at a national scale. Biol. Conserv. 132, 481–489 (2006).

7. Hall, D. M. et al. The city as a refuge for insect pollinators. Conserv. Biol. 31, 24–29 (2016).

8. Baldock, K. C. R. et al. A systems approach reveals urban pollinator hotspots and conservation opportunities. Nat. Ecol. Evol. 3, 1–15 (2019).

9. Samuelson, A. E., Gill, R. J., Brown, M. J. F. F. & Leadbeater, E. Lower bumblebee colony reproductive success in agricultural compared with urban environments. Proc. R. Soc. B Biol. Sci. 285, 20180807 (2018).

10. Lecocq, A., Kryger, P., Vejsnæs, F. & Bruun Jensen, A. Weight Watching and the Effect of Landscape on Honeybee Colony Productivity: Investigating the Value of Colony Weight Monitoring for the Beekeeping Industry. PLoS One 10, e0132473 (2015).

11. Plascencia, M. & Philpott, S. M. Floral abundance, richness, and spatial distribution drive urban garden bee communities. Bull. Entomol. Res. 107, 658–667 (2017).

12. Seeley, T. D. The wisdom of the hive : the social physiology of honey bee colonies. (Harvard University Press, 1995).

13. Thomson, D. Competitive interactions between the invasive European honey bee and native bumblebees. Ecology 85, 458–470 (2004).

14. von Frisch, K. The Dance Language and Orientation of Bees. (Belknap Press of Harvard University Press, 1967).

15. Couvillon, M. J. & Ratnieks, F. L. W. Environmental consultancy : dancing bee bioindicators to evaluate landscape “health”. Front. Ecol. Evol. 3, 1–8 (2015).

16. Couvillon, M. J., Schürch, R. & Ratnieks, F. L. W. Waggle Dance Distances as Integrative Indicators of Seasonal Foraging Challenges. PLoS One 9, e93495 (2014).

17. Waddington, K. D. et al. Comparisons of forager distributions from matched in suburban honey bee colonies environments. Behav. Ecol. Sociobiol. 35, 423–429 (1994).

18. Visscher, P. K. & Seeley, T. D. Foraging Strategy of Honeybee Colonies in a Temperate Deciduous Forest. Ecology 63, 1790 (1982).

19. Requier, F. et al. Honey bee diet in intensive farmland habitats reveals an unexpectedly high flower richness and a major role of weeds. Ecol. Appl. 25, 881–890 (2015).

20. Hung, K.-L. J., Kingston, J. M., Albrecht, M., Holway, D. A. & Kohn, J. R. The worldwide importance of honey bees as pollinators in natural habitats. Proc. R. Soc. B Biol. Sci. 285, 20172140 (2018).

21. Goulson, D. Bumblebees : behaviour, ecology, and conservation. (Oxford University Press, 2010).

22. Osborne, J. L. et al. Quantifying and comparing bumblebee nest densities in gardens and countryside habitats. J. Appl. Ecol. 45, 784–792 (2008).

23. Kabisch, N., Strohbach, M., Haase, D. & Kronenberg, J. Urban green space availability in European cities. Ecol. Indic. 70, 586–596 (2016).

24. Williams, N. M. et al. Native wildflower plantings support wild bee abundance and diversity in agricultural landscapes across the United States. Ecol. Appl. 25, 2119–2131 (2015).

25. Schürch, R. et al. Dismantling Babel: creation of a universal calibration for honey bee waggle dance decoding. Anim. Behav. 150, 139–145 (2019).

26. Couvillon, M. J. et al. Intra-dance variation among waggle runs and the design of efficient protocols for honey bee dance decoding. Biol. Open 1, 467–472 (2012).

27. Graham, M. H. Confronting Multicollinearity in Ecological Multiple Regression. Ecology 84, 2809–2815 (2003).

28. Samuelson, A. E. & Leadbeater, E. A land classification protocol for pollinator ecology research: An urbanization case study. Ecol. Evol. 8, 5598–5610 (2018).

29. Schürch, R. et al. Incorporating variability in honey bee waggle dance decoding improves the mapping of communicated resource locations. 1143–1152 (2013). doi: 10.1007/s00359-013-0860-4

30. Barton, K. MuMIn: Multi-Model Inference. (2018).

31. Agostinelli, C. & Lund, U. R package ‘circular’: Circular Statistics (version 0.4-93). (2017).

32. Hajage, D. ascii: Export R objects to several markup languages. (2011).

33. Hijmans, R. J. raster: Geographic Data Analysis and Modeling. (2017).

34. Bivand, R. S., Pebesma, E. & Gomez-Rubio, V. Applied spatial data analysis with {R}, Second edition. (Springer, NY, 2013).

35. Bivand, R., Keitt, T. & Rowlingson, B. rgdal: Bindings for the ‘Geospatial’ Data Abstraction Library. (2018).

36. Bivand, R. & Rundel, C. rgeos: Interface to Geometry Engine – Open Source (‘GEOS’). (2017).

37. Ogle, D. H. FSA: Fisheries Stock Analysis. (2018).

38. Bates, D., Mächler, M., Bolker, B. & Walker, S. Fitting Linear Mixed-Effects Models Using {lme4}. J. Stat. Softw. 67, 1–48 (2015).

39. Eklund, A. beeswarm: The Bee Swarm Plot, an Alternative to Stripchart. (2016).

40. Harrell Jr, F. E. & Dupont, C. Hmisc: Harrell Miscellaneous. (2018).

