## Supplementary figures for "Dancing bees evaluate agricultural forage resources as inferior to central urban land"

### Supplementary Information

Ash E. Samuelson[1*], Roger Schürch[2] & Ellouise Leadbeater[1]

[1] School of Biological Sciences, Royal Holloway University of London, Egham, United Kingdom

[2] Department of Entomology, Virginia Tech, Blacksburg, VA, USA

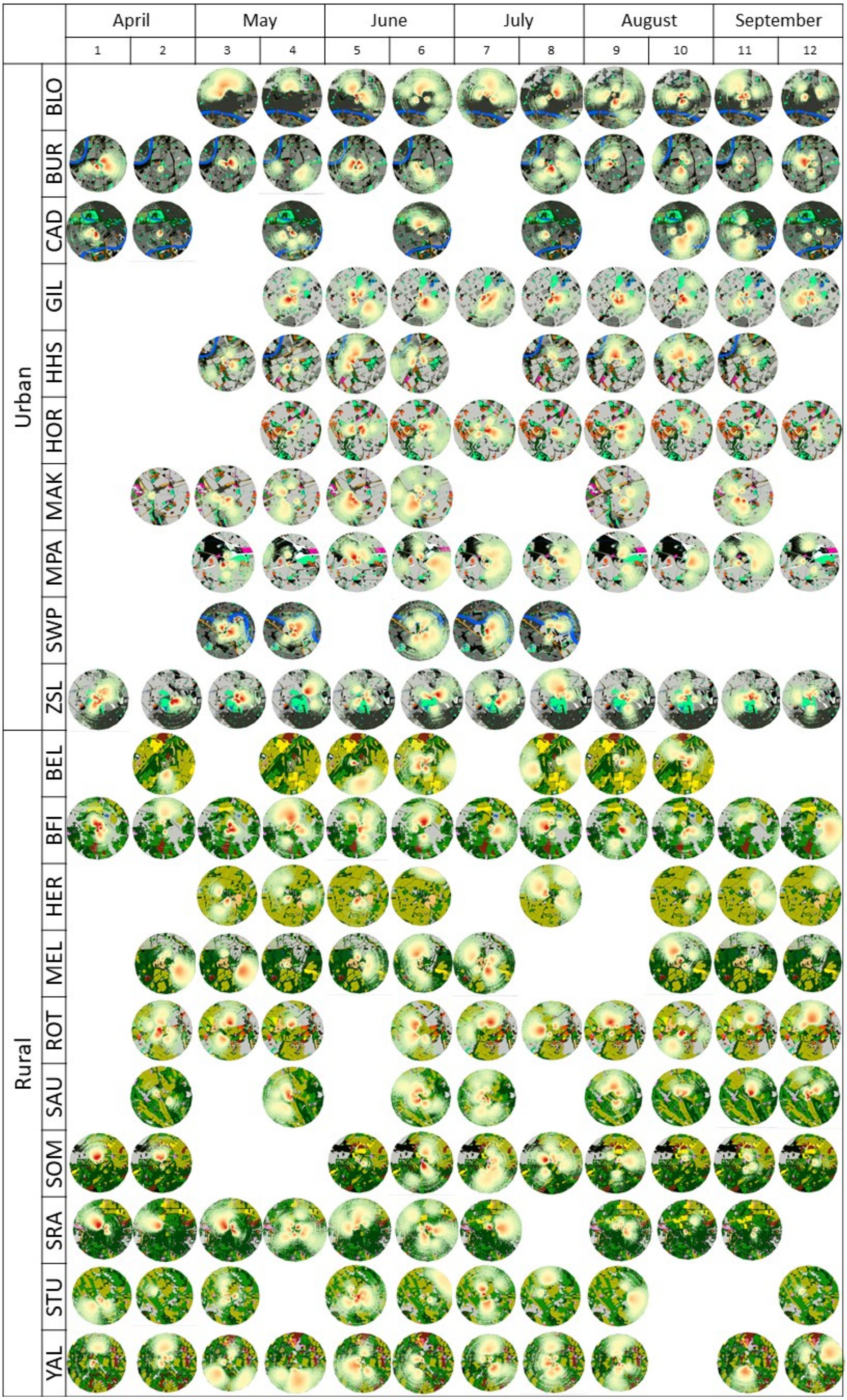
April
May
June
July
August
September
1
2
3
4
5
6
7
8
9
10
11
12
Urban
BLO
BUR
CAD
GIL
HHS
HOR
MAK
MPA
SWP
ZSL
Rural
BEL
BFI
HER
MEL
ROT
SAU
SOM
SRA
STU
YAL

(previous page) **Figure S1.** Waggle dance plots from ten urban sites and ten rural sites over six months (12 fortnightly timepoints). Each circle shows the dances recorded on a single filming period (up to 3 hours) at a single location. Waggle dances are displayed as probability heatmaps generated from 1000 simulations of each dance allowing incorporation of variability in distance and angle communication (Schürch *et al.*, 2013). Dance plots are overlaid on GIS land-use maps (radius 2500m) produced for land-use preference analysis.

**Figure S2.** Beeswarm plots of log-transformed waggle run durations for a subset of two urban (blue) and two rural (green) sites at which dances were recorded in both a) 2016 and b) 2017. Black lines indicate median values.

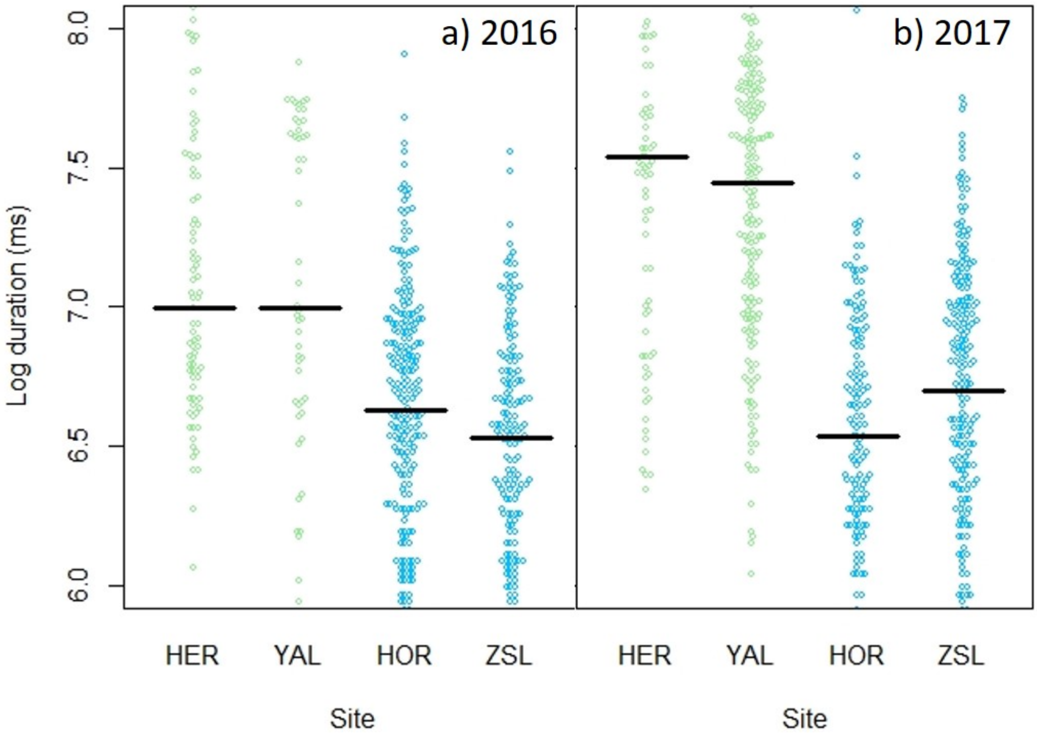
